# Islet architecture in adult mice is actively maintained by Robo2 expression in β cells

**DOI:** 10.1101/2022.09.07.506980

**Authors:** Bayley J. Waters, Barak Blum

## Abstract

Pancreatic islets of Langerhans have a non-random spatial architecture, which is important to proper glucose homeostasis. Islet architecture forms during embryonic development, in a morphogenesis process partially involving expression of Roundabout (Robo) receptors in β cells, and their ligand, Slit, in the surrounding mesenchyme. Whether islet architecture is set during development and remains passive in adulthood, or whether it requires active maintenance throughout life, has not been determined. To distinguish between these two models, we conditionally deleted *Robo2* in β cells of adult mice and observed their islet architecture following a two-month chase. Here, we show that deleting *Robo2* in adult β cells causes significant loss of islet architecture without affecting β cell identity, maturation, or stress, indicating that Robo2 plays a role in actively maintaining adult islet architecture. Understanding the factors required to maintain islet architecture, and thus optimize islet function, is important for developing future diabetes therapies.

## Introduction

The spatial arrangement of endocrine cell types within the islets of Langerhans is important to their function of regulating blood glucose. β cells in the islet function optimally when in contact with other β cells, whereby they can electrically couple to coordinate insulin secretion (Brereton et al., 2006; Farnsworth et al., 2014). Both humans and mice exhibit islet architecture which prioritizes β cell-β cell homotypic interactions (Hoang et al., 2014), with humans showing more complex islet architecture in the form of β cell clusters interspersed with α, δ, and PP cells (Bonner-Weir et al., 2015), and mice showing a more uniform β cell core surrounded by a mantle of other endocrine cell types.

Loss of islet architecture is seen in diabetes mellitus in both humans and mice (Brereton et al., 2015; Steiner et al., 2010), but it is unclear whether architecture loss is an attempted adaptation to, or a pathological result of, increased diabetogenic stress (Adams and Blum, 2022). Moreover, evidence suggests that loss of synchronous insulin secretion among β cells contributes to blunted insulin sensitivity over time, potentially exacerbating dysglycemia (Matveyenko et al., 2012; Satin et al., 2015). Thus, understanding factors involved in the formation and maintenance of islet architecture is a key consideration in therapies aimed at improving or replacing native islet function.

Islet architecture is established during development, when endocrine progenitor cells migrate away from the pancreatic ducts and cluster together, simultaneously beginning the differentiation process to achieve their mature endocrine identities (Pan and Wright, 2011; Shih et al., 2013; Sznurkowska et al., 2020). We previously reported that the axon guidance molecules Roundabout (Robo) receptors are required to establish islet architecture in development (Adams et al., 2018). Deletion of *Robo2* in the β cells of mice during development results in loss of classic murine core-mantle islet architecture, but does not affect β cell identity, maturation, or stress (**Figure 1A**). This aberrant islet architecture exhibits significant loss of homotypic β cell-β cell interactions, and reduced synchroneity of β cell insulin pulses (Adams et al., 2021). These data indicate that failure to establish correct islet architecture during development has lifelong consequences on islet function. However, whether normal islet architecture remains in place once established, or requires continued cues to be maintained, has not been determined.

**Figure 1:**
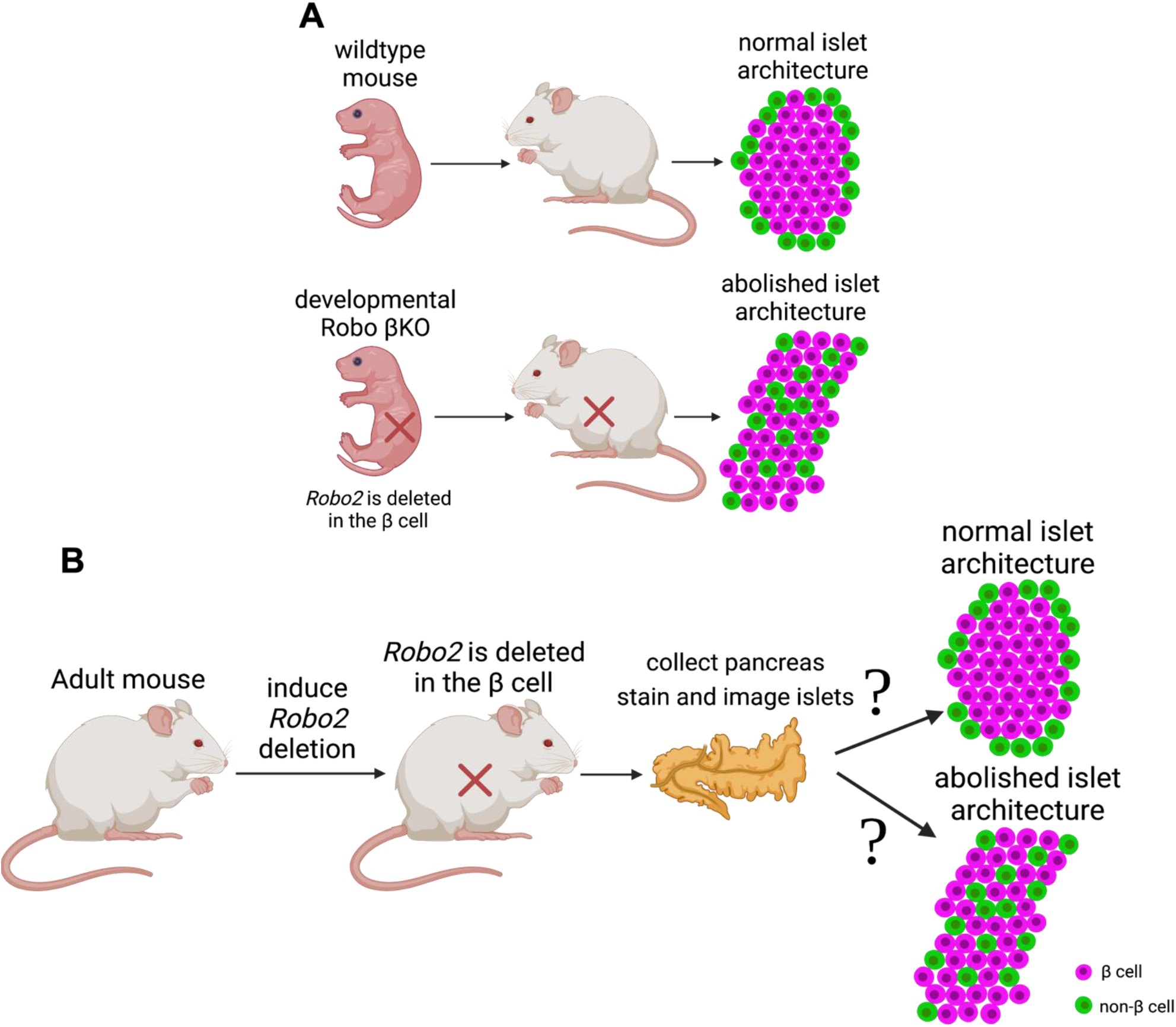
Is islet architecture actively maintained in adulthood? **(A)** Deletion of *Robo2* in the β cells during development results in total loss of islet architecture. **(B)** Possible outcomes following *Robo2* deletion from the β cells of adult mice. If islet architecture is passive in the adult and does not require maintenance by Robo2, the islet will have normal architecture following *Robo2* deletion (top). If Robo2 is required to actively maintain islet architecture, the islet will exhibit cell type intermixing (bottom).

In the current study, using temporally-controlled *Robo2* deletion as a model, we followed up on Robo’s guidance role in islet development to assess whether *Robo2* may also be involved in continued maintenance of the adult islet. Here, we report that islet architecture is actively maintained in adulthood, and that *Robo2* is required for this maintenance.

## Results

We aimed to explore a fundamental question in islet development: whether islets of Langerhans, whose tissue architecture is carefully orchestrated during development, require continued maintenance after architecture is established. We have previously shown that expression of Robo in β cells is required for endocrine cell type sorting as mouse islets form during development (Adams *et al*., 2018). However, Robo continues to be expressed in the adult islet, and is strongly downregulated in early stages of obesity, concomitant with compensatory islet expansion (Adams *et al*., 2018). We hypothesized that Robo is also required for active maintenance of islet architecture in adulthood. To test this hypothesis, we deleted *Robo2* in the β cells of adult mice using an inducible Cre-Lox system. **Figure 1B** illustrates two possible outcomes following *Robo2* β cell-deletion. If islet architecture does not require active maintenance after being established, then deletion of *Robo2* in adulthood will not impact islet architecture (**top**). If islet architecture does require active maintenance, and Robo2 continues to play a similar role in positional cell guidance in adulthood as it does in development, then intra-islet endocrine cell type admixture is expected following *Robo2* deletion in adults (**bottom**).

To delete *Robo2* in the β cells of mice after they reached adulthood, we used a β cell-specific, estrogen receptor-dependent Cre driver (*Ins1-CreERT2*) to delete the floxed portion of the *Robo2* allele (**Supplemental Figure 1A**). We administered tamoxifen (TAM) in corn oil via oral gavage (OG) to mice aged 10-29 weeks. Mice received 5 consecutive days of TAM dosing, followed by 8 weeks of rest to allow for reduced Robo2 protein expression and any potential cell movement, before their pancreata were collected for analysis (**Figure 2A**). Study mice also possessed a copy of nuclear fluorescent reporter construct Rosa26^lox-stop-lox-H2b-mCherry^, which provided an indication of the success of our Cre induction and thus *Robo2* deletion, as β cell nuclei fluoresced mCherry once the associated stop codon was deleted (**Supplemental Figure 1A,B**).

**Figure 2:**
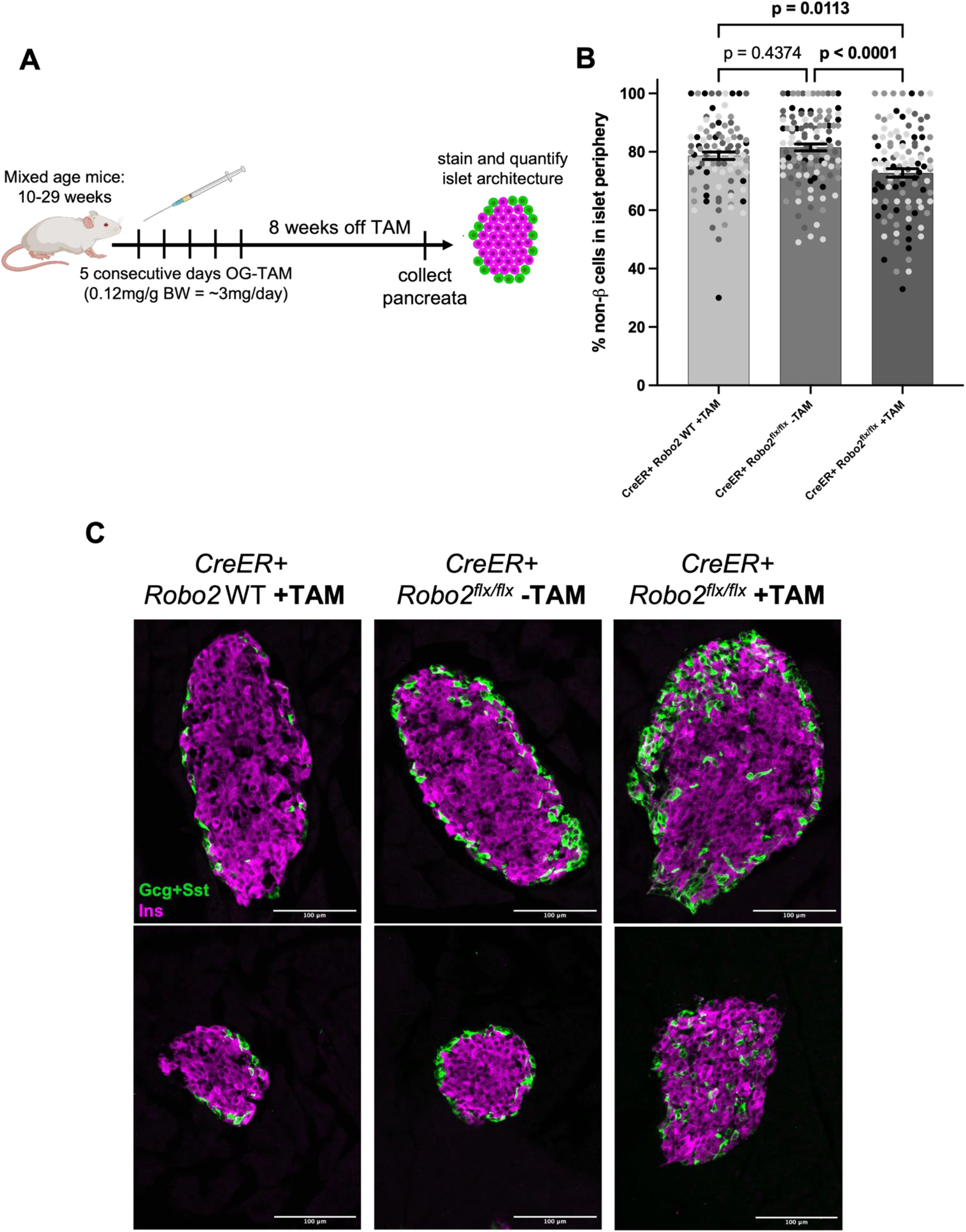
Deleting Robo2 in adult mouse β cells causes significant loss of islet architecture. **(A)** Mice were enrolled to receive tamoxifen (TAM) or vehicle (corn oil) for 5 consecutive days at a dose of 0.12mg/g body weight. 8 weeks following the final dose, mice were sacrificed and pancreata collected for analysis of islet architecture. **(B)** Quantification of the percentage of mantle-localized non-β (α and δ) cells over the total number of non-β cells in the islet. Each color represents different islets from the same mouse. Adult mice with *Robo2* deleted (right bar) show a significantly lower percentage of mantle-localized non-β cells in comparison to controls, indicating disrupted islet architecture. N=4 mice per group, 17-37 islet per mouse, analyzed via 2-way ANOVA with Bonferroni post-hoc test for multiple comparisons. **(C)** Representative images of islet architecture in *Robo2^+/+^* (left), *Robo2*^flx/flx^ mice without TAM induction (middle), and *Robo2*^flx/flx^ mice with TAM induction (right). Insulin in magenta, glucagon and somatostatin in green.

For quantification of islet architecture, non-β (α and δ) cells were visualized and counted together. For each islet, quantification of the percentage of non-β cells in the islet mantle indicates the relative admixture of islet architecture, with lower percentages indicating more severe admixture. A standard wildtype mouse islet will have nearly all non-β cells in the islet mantle, approaching a value of 100%.

We found that deletion of *Robo2* in adulthood resulted in a significant reduction in the number of non-β cells in the mantle, indicating a notable loss of normal islet architecture (**Figure 2B,C**). *Ins1-CreERT2^Tg/0^; Robo2^flx/flx^* mice which received TAM (*CreER+ Robo2^flx/flx^* +TAM) had significantly more intermixed islet architecture than both *Ins1-CreERT2^Tg/0^; Robo2^+/+^* mice who received TAM (*CreER+* WT +TAM, p=0.0113), and *Ins1-CreERT2^Tg/0^; Robo2^flx/flx^* which did not receive TAM (*CreER+ Robo2^flx/flx^* −TAM, p<0.0001). Thus, our results indicate that Robo2 is required to maintain islet architecture in adult mice, and more broadly, that islet architecture is not simply “set” during development but requires active maintenance after development is complete.

Disruptions in islet architecture are seen in all forms of diabetes mellitus, including in human patients with diabetes and mouse models of diabetes (Brereton *et al*., 2015; Steiner *et al*., 2010). Diabetes also presents with loss of β cell maturity (Nimkulrat et al., 2021), which may contribute to the observed changes in islet architecture. To confirm that architecture changes in our model were not due to loss of β cell maturity, we stained for β cell maturity markers urocortin 3 (Ucn3) and MafA. We observed no notable differences between the experimental group and CreER+ wildtype controls in overall expression of Ucn3 and MafA (**Figure 3A**), indicating that changes in islet architecture are not due to changes in β cell maturity.

**Figure 3:**
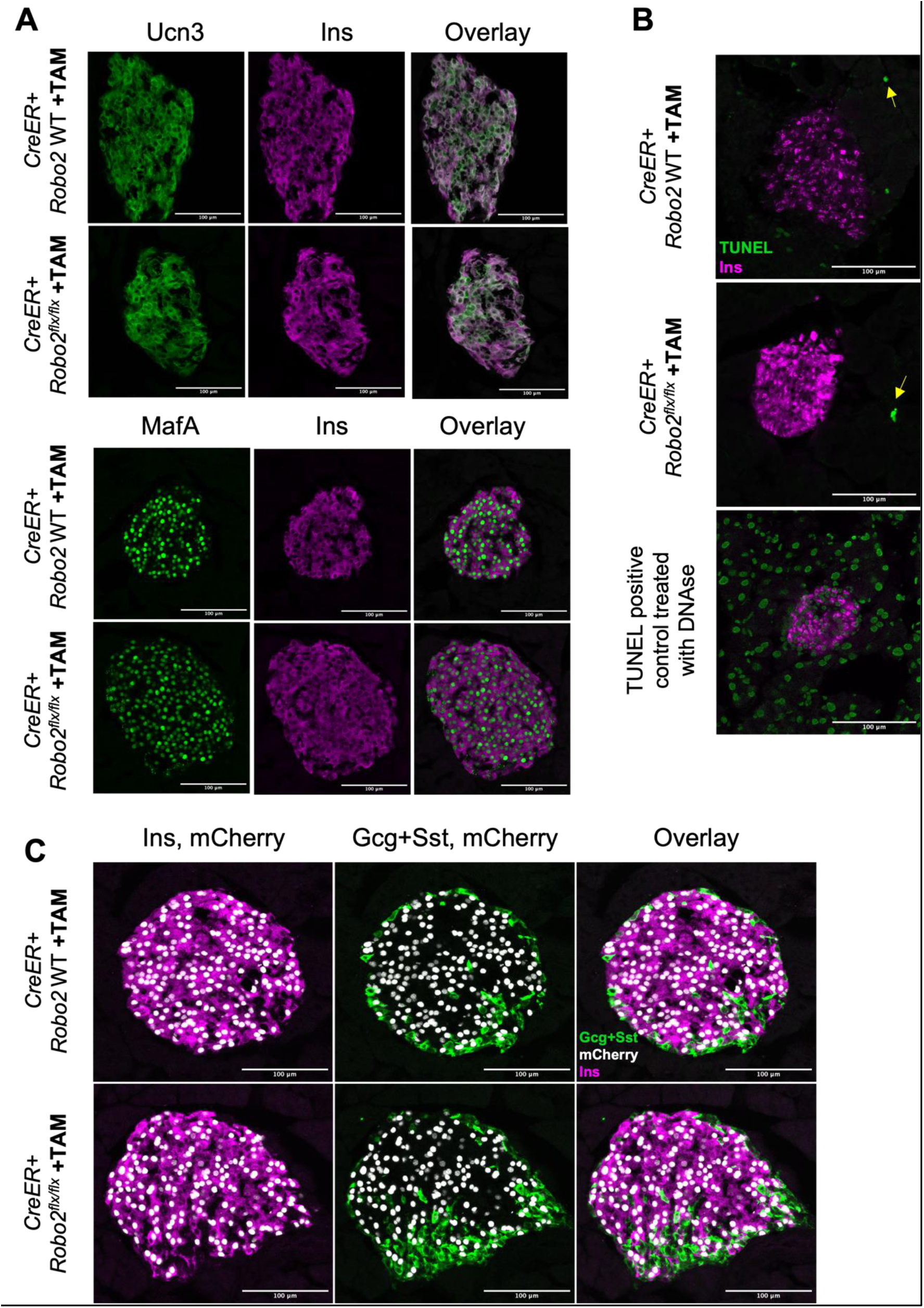
Loss of islet architecture in adult Robo2 mutant mice is not due to loss of ß cell identity or maturity, or β cell death. **(A)** Islets from *Robo^+/+^* and TAM-induced *Robo2*^flx/flx^ mice show similar levels of Ucn3 and MafA. **(B)** Islets from *Robo^+/+^* and TAM-induced *Robo2*^flx/flx^ mice do not have significant differences in cell death (TUNEL). Yellow arrows indicate presence TUNEL staining in cell nuclei. **(C)** Changes in islet architecture are not due to β cell transdifferentiation, as β cell lineage tracer mCherry localizes only in β cells and not in non-β cells. Insulin in magenta, glucagon and somatostatin in green, mCherry in white.

Similarly, the observed changes in adult islet architecture could not be explained by differences in cell death, as number of TUNEL-positive cells were negligible, and not different among groups (**Figure 3B**). Presence of *Ins1-CreERT2-driven* mCherry only in β cell nuclei, and not in α or δ cell nuclei, indicates that changes in architecture are not due to β cell transdifferentiation into other endocrine cell types (**Figure 3C**). Thus, aberrations in islet architecture observed following *Robo2* deletion in adulthood appear to result directly from loss of Robo2, and not from β cell dedifferentiation, transdifferentiation, or death, as have been reported in development (Adams *et al*., 2018).

## Discussion

We tested the hypothesis that islet architecture still requires active maintenance even after the islet has reached its adult state, by using an inducible model of *Robo2* β cell deletion. Here, we show that islet architecture requires active signaling maintenance after development is complete. This finding is notable, as it has not been shown to date that islets require continued maintenance cues after they achieve mature architecture. Additionally, our discovery that Robo2 is necessary to maintain islet architecture in adulthood indicates the potency of Robo2 as a guidance molecule in the mature islet, since reduction in this receptor alone caused significant cell movement and loss of mantle-core architecture despite the presumed presence of all other regulatory cues in the islet.

This work reveals that not only does Robo play a role in the adult islet, but also controls islet architecture under homeostatic conditions, wherein there is no demand on the islet to expand and proliferate. Robo is required in endocrine progenitors and embryonic β cells to establish islet architecture as the islet forms throughout development (Adams *et al*., 2018), yet the islet continues to expand after β cells reach maturity. Similarly, islets must expand in response enhanced demand for insulin, such as in pregnancy or under obesogenic stimulus (Rieck and Kaestner, 2010; Weir and Bonner-Weir, 2004). In all cases where islets expand, whether in development or under certain metabolic conditions, islet architecture must remain consistent to preserve optimal islet function. Indeed, it is hypothesized that the disrupted islet architecture in diabetes represents a collapse of normal architecture as islets fail to remodel under expansion-inducing conditions (Adams and Blum, 2022). Alternatively, loss of islet architecture in diabetes may represent an attempt by islets to “loosen” their structure in order to make remodeling easier in the face of increased insulin demand (Adams and Blum, 2022). In either case, it appears that islets must receive cues to regulate their architecture during proliferation events. It would be interesting to study the role that Robo may play in regulating islet architecture during islet expansion.

Increasingly relevant is the importance of islet architecture in preserving the islet function of donor islets, or in building functional islets from stem cells. Islets in culture are observed to lose their architecture and regain it when transplanted (Lavallard et al., 2016). Similarly, islets derived from stem cells function better when induced to form mature architecture (Balboa et al., 2022). Research continues to explore the essential role of extracellular matrix (ECM) proteins in regulating islet architecture, including supplementing Matrigel with key ECM factors (Tixi et al., 2022; Tremmel et al., 2022). Slit is the canonical ligand for Robo receptors, and has also been shown to impact islet formation in developing mice (Gilbert et al., 2020). Slit from the pancreatic mesenchyme during development may thus interact with Robo on the developing β cell to direct endocrine cell type sorting (Waters and Blum, 2022). It is possible that Slit from peri-islet sources may also interact with Robo on the adult β cell to maintain endocrine cell arrangement in adulthood. Further elucidation of the mechanisms behind Robo’s role in maintaining adult islet architecture, and the source of the ligands by which its function is regulated, may play a key part in maintaining healthy islet architecture in islets intended for transplant.

## Methods

### Animals

The experimental protocol for animal usage was reviewed and approved by the University of Wisconsin-Madison Institutional Animal Care and Use Committee (IACUC) under Protocol #M005221, and all animal experiments were conducted in accordance with the University of Wisconsin-Madison IACUC guidelines under the approved protocol. All mouse strains were maintained on a mixed genetic background. *Ins1-CreERT2* (Tamarina et al., 2014), *Robo2*^flx^ (Lu et al., 2007), and *Rosa26*^lox-stop-lox-H2B-mCherry^ (Blum et al., 2014) mice were previously described. Control colony mates in all analyses were either Cre+ Robo2^+/+^ or Cre+ Robo2^flx/flx^ without tamoxifen.

### Tamoxifen solution and dosing

Tamoxifen solution was prepared as previously described (Donocoff et al., 2020). In brief, 300mg powdered tamoxifen (Sigma, T5648) was dissolved in 10ml of syringe-filtered corn oil at 50°C for 2 hours, and distributed into 1mL aliquots. Both tamoxifen solution and vehicle solution were stored at 4°C in a light-protected box for the duration of the dosing week. TAM and vehicle were preheated at 50°C for 30 minutes before filling syringes each day. Mice were weighed on the first day of dosing, and received 0.12mg/g of this body weight of tamoxifen each day for five days by oral gavage. Mice were sacrificed eight weeks after the last tamoxifen dose for tissue collection.

### Tissue collection and sectioning

Mice were euthanized with CO2 according to standard protocol and their pancreata immediately collected for fixation. Pancreata were fixed in 4% PFA in PBS for 3 hours at 4°C. Samples were then washed, equilibrated in 30% sucrose in PBS, and embedded in FSC 22 Blue. Tissue was cryosectioned with four 10um slices per slide, each 400um apart.

### Immunostaining and confocal imaging

Slides were stained using a standard immunofluorescence protocol. The following primary antibodies were used: guinea pig anti-insulin 1:6 (Agilent, IR002), rabbit anti-glucagon 1:200 (Cell Signaling Technology, 2760S), rabbit anti-somatostatin 1:500 (Phoenix Pharmaceuticals, G-060-03), rabbit anti-Ucn3 (Phoenix Pharmaceuticals, H-019-29), and rabbit anti-MafA (Cell Signaling Technology, 79737S). The following secondary antibodies were used: donkey anti-guinea pig AF647 (Jackson, 706-605-148), donkey anti-rabbit AF488 (Invitrogen, A21206), and DAPI. TUNEL staining was performed following citrate buffer antigen retrieval using an *in situ* cell death detection kit (Roche, 11684795910). Positive control slide was treated with deoxyribonuclease to induce DNA damage (Sigma, D4263). Slides were imaged using a Nikon Upright FN1 provided by the University of Wisconsin Optical Imaging Core (UWOIC, grant #1S10OD025040-01).

### Analysis and statistics

Cells were counted from sum intensity projected Z-stack images of islets, comprised of seven 1um slices, using the Cell Counter tool on FIJI (Fiji Is Just ImageJ). Cells were counted as previously described (Adams *et al*., 2018). Groups were compared using a 2-way ANOVA with Bonferroni post-hoc analysis for multiple comparisons. Data are presented as average ±SEM. All statistics were done using Prism GraphPad. P-values below 0.05 were considered significant.

## Data availability

All data generated during and/or analyzed during the current study are available from the corresponding author on reasonable request.

## Acknowledgements

We would like to thank members of the Blum lab, particularly Dr. Melissa Adams, Dex Nimkulrat, and Cyrus Sethna for fruitful discussion, technical advice, and manuscript edits, and to Debayan De Bakshi for manuscript edits and support. We thank the University of Wisconsin Optical Imaging Core managing director Lance Rodenkirch for microscopy support. Parts of some figures were created with biorender.com.

## Author contributions

BB and BJW conceived and designed experiments. BJW performed experiments and analyzed data. BB and BJW wrote and edited the manuscript.

## Funding

NIH R01DK121706, T32 HD041921, University of Wisconsin-Madison Optical Imaging Core (UWOIC) support grant 1S10OD025040-01

## Declaration of interest

The authors declare no conflict of interests.

## Supplemental Figure 1

**Supplemental Figure 1:**
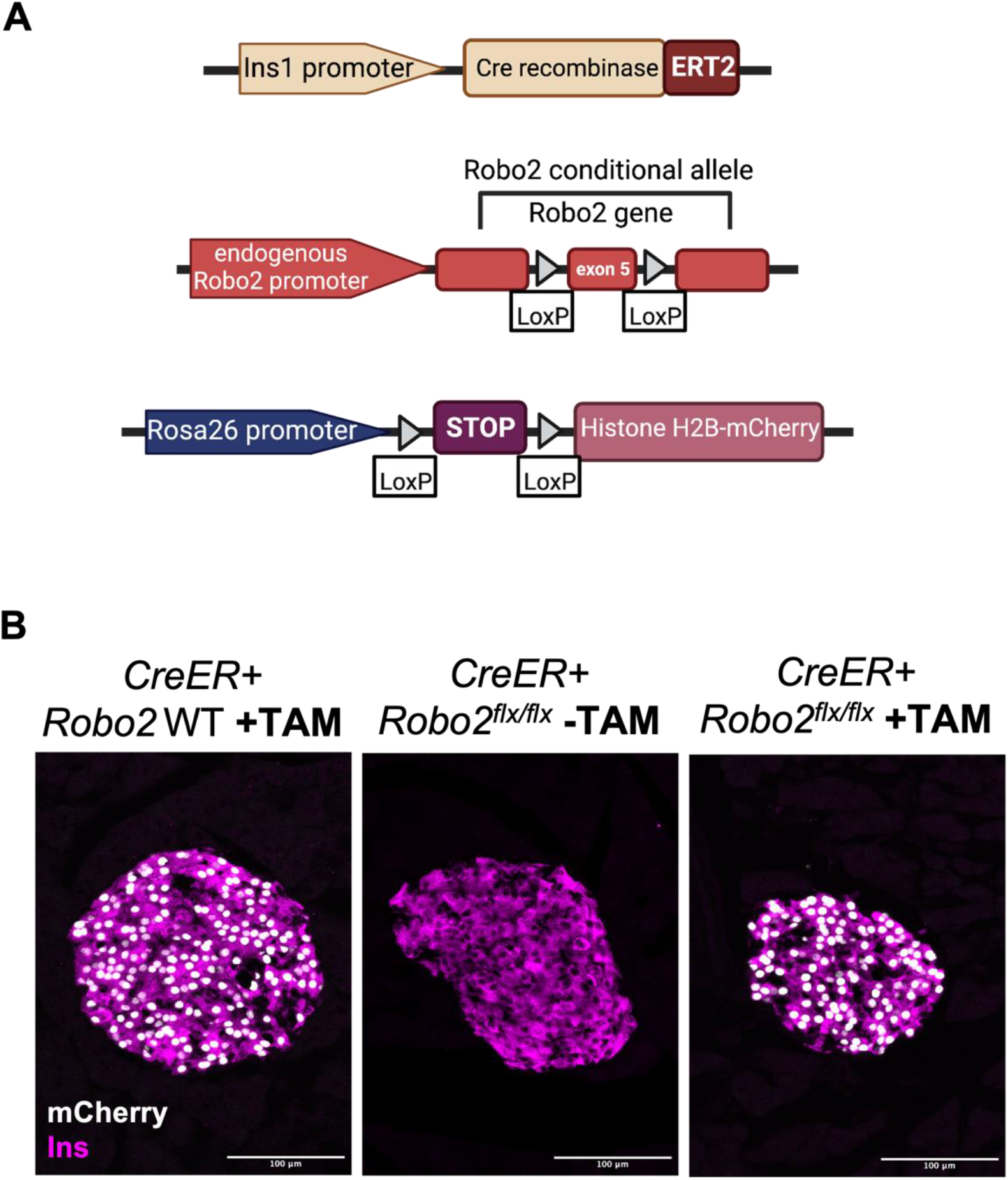
**(A)** Genetic constructs used in study mice. **(B)** Presence of nuclear fluorescent reporter mCherry in the β cell following TAM dosing indicates that the Cre-Lox system has high penetrance and that *Robo2* has been deleted in these cells. The control group that did not receive TAM does not express mCherry (middle).

## Notes

### Competing Interest Statement

The authors have declared no competing interest.

